# Mito-FUNCAT-FACS reveals cellular heterogeneity in mitochondrial translation

**DOI:** 10.1101/2022.01.03.474764

**Authors:** Yusuke Kimura, Hironori Saito, Tatsuya Osaki, Yasuhiro Ikegami, Taisei Wakigawa, Yoshiho Ikeuchi, Shintaro Iwasaki

**Affiliations:** Department of Computational Biology and Medical Sciences, Graduate School of Frontier Sciences, The University of Tokyo, Kashiwa, Chiba 277-8561, Japan; RNA Systems Biochemistry Laboratory, RIKEN Cluster for Pioneering Research, Wako, Saitama 351-0198, Japan; Institute of Industrial Science, The University of Tokyo, Meguro-ku, Tokyo 153-8505, Japan; Institute for AI and Beyond, The University of Tokyo, Bunkyo-ku, Tokyo 113-0033, Japan

## Abstract

Mitochondria possess their own genome that encodes components of oxidative phosphorylation (OXPHOS) complexes, and mitochondrial ribosomes within the organelle translate the mRNAs expressed from mitochondrial genome. Given the differential OXPHOS activity observed in diverse cell types, cell growth conditions, and other circumstances, cellular heterogeneity in mitochondrial translation can be expected. Although individual protein products translated in mitochondria have been monitored, the lack of techniques that address the variation in overall mitochondrial protein synthesis in cell populations poses analytic challenges. Here, we adapted mitochondrial-specific fluorescent noncanonical amino acid tagging (FUNCAT) for use with fluorescence-activated cell sorting (FACS) and developed mito-FUNCAT-FACS. The click chemistry-compatible methionine analog L-homopropargylglycine (HPG) enabled the metabolic labeling of newly synthesized proteins. In the presence of cytosolic translation inhibitors, HPG was selectively incorporated into mitochondrial nascent proteins and conjugated to fluorophores via the click reaction (mito-FUNCAT). The application of *in situ* mito-FUNCAT to flow cytometry allowed us to disentangle changes in net mitochondrial translation activity from those of the organelle mass and detect variations in mitochondrial translation in cancer cells. Our approach provides a useful methodology for examining mitochondrial protein synthesis in individual cells.

## Introduction

Mitochondria are the major suppliers of cellular ATP via oxidative phosphorylation (OXPHOS). As a consequence of the symbiosis of a bacterial ancestor, mitochondrion still maintains its own genome and expresses its own genes. In addition to noncoding RNAs (2 rRNAs and 22 tRNAs), mammalian mitochondrial DNA (mtDNA) encodes 13 mRNAs that are essential components of OXPHOS complexes (Anderson et al. 1981). Impairment of mitochondrial translation, such as dysfunction of mitochondrial tRNAs, often causes OXPHOS dysfunction and leads to diseases (Scharfe et al. 2009; De Silva et al. 2015; Suomalainen and Battersby 2018; Webb et al. 2020), including mitochondrial myopathy, encephalopathy, lactic acidosis, and stroke-like episodes (MELAS) and myoclonic epilepsy with ragged-red fibers (MERRF) (Yasukawa et al. 2000a, 2000b; Kirino et al. 2005; Rötig 2011; Keilland et al. 2016; Asano et al. 2018; Morscher et al. 2018).

Mitochondrial protein synthesis is driven by 55S mitochondrial ribosomes (or mitoribosomes). Owing to the recent emergence of structures at near-atomic resolutions, the architecture of mitoribosomes has been revealed to be distinct from bacterial and eukaryotic cytoplasmic ribosomes (or cytoribosomes) (Kummer and Ban 2021). *In organello* translation has been investigated through various approaches, such as the following: 1) sucrose density gradient ultracentrifugation for polysome isolation and subsequent mRNA quantification via quantitative reverse transcription-PCR (RT-qPCR) or Northern blot (Fung et al. 2013; Antonicka et al. 2013; Zhang et al. 2014; Grimes et al. 2014; Pearce et al. 2017; Cahoon and Qureshi 2018); 2) ribosome profiling (Iwasaki et al. 2016; Suzuki et al. 2020; Kashiwagi et al. 2021), a method based on RNase footprinting by ribosomes and following deep sequencing (Ingolia et al. 2009), and the derivative methods tailored for mitochondrial translation (Rooijers et al. 2013; Couvillion et al. 2016; Couvillion and Churchman 2017; Gao et al. 2017; Pearce et al. 2017; Morscher et al. 2018; Li et al. 2021; Schöller et al. 2021); 3) pulse stable isotope labeling by amino acids in cell culture (pSILAC) following mass spectrometry (Imami et al. 2021); 4) *in situ* cryo-electron tomography (Pfeffer et al. 2015; Englmeier et al. 2017); 5) *in vitro* reconstitution with purified factors (Lee et al. 2021); and 6) *in vitro* silencing of translation in purified mitochondria (Cruz-Zaragoza et al. 2021).

The most conventional method used by mitochondrial protein synthesis studies is to label translation products with radioactive amino acids (*e.g.*, L-[^35^S]-methionine). The relatively large pool of cytoribosomes and translation products hampers the detection of proteins synthesized in mitochondria. Thus, cells are treated with cytoribosome-specific inhibitors such as cycloheximide (CHX), anisomycin (ANS), and emetine (EME) (Garreau de Loubresse et al. 2014; Wong et al. 2014) to suppress cytosolic translation, allowing the specific labeling of mitoribosome-synthesizing proteins (Jeffreys and Craig 1976; Weraarpachai et al. 2009).

Despite the high sensitivity of the radioactive labeling of nascent proteins, the availability of alternative methods involving fluorescence is also an advantage (regardless of whether translation occurs in the cytoplasm or mitochondria) since these alternative methods are applicable to microscopy and flow cytometry analyses (Iwasaki and Ingolia 2017). The metabolic incorporation of methionine analogs with click-reactive moieties, such as L-homopropargylglycine (HPG) and L-azidohomoalanine (AHA) (Supplemental Fig. S1A), allows the conjugation of fluorophores to proteins synthesized during a given period. This technique, known as fluorescent noncanonical amino acid tagging (FUNCAT), is employed to measure cytosolic translation on sodium dodecyl sulfate-polyacrylamide gel electrophoresis (SDS-PAGE) (Yoon et al. 2012), under a microscope (Beatty et al. 2006; Beatty and Tirrell 2008; Roche et al. 2009; Dieterich et al. 2010; Tcherkezian et al. 2010; Hinz et al. 2012; Yoon et al. 2012), and through flow cytometry (Beatty and Tirrell 2008; Signer et al. 2014).

In contrast to cytosolic translation, the application of FUNCAT to mitochondrial translation (mito-FUNCAT) has been restricted to bulk on-gel assays (Zhang et al. 2014) (on-gel mito-FUNCAT) and microscopy analyses (Estell et al. 2017; Yousefi et al. 2021; Zorkau et al. 2021) (*in situ* mito-FUNCAT) (Supplemental Fig. S1B). However, these approaches (and the other methods mentioned above) may miss a subpopulation of cells because bulk on-gel assays average all cells and microscopy covers only a limited number of cells. In this work, we developed the mito-FUNCAT-fluorescence-activated cell sorting (FACS) method for the high-throughput quantification of mitochondrial protein synthesis in cells in response to biogenesis and functional activation of mitochondria and investigated the cellular heterogeneity of *in organello* translation.

## Results and discussion

### In-cell visualization of mitochondrial protein synthesis

To investigate mitochondrial translation, we metabolically labeled newly synthesized proteins generated within mitochondria with the click-reactive methionine analogs HPG and AHA (Supplemental Fig. S1A). Although these analogs can be incorporated into both cytosolic and mitochondrial nascent peptides, halting cytosolic translation with the cytoribosome-specific inhibitor ANS (Garreau de Loubresse et al. 2014) resulted in only active mitochondrial translation over a given period of analog incubation (Fig. 1A). The resultant HPG-labeled nascent proteins were conjugated with fluorophores via an *in vitro* click reaction and visualized via SDS-PAGE (on-gel mito-FUNCAT) (Fig. 1B and Supplemental Fig. S1B). Essentially, the same pattern of signals could be obtained with other cytosolic translation inhibitors, CHX and EME (Supplemental Fig. S1C). The use of this method was further validated for the assessment of mitochondrial protein synthesis by the recovery of “HPGylated” proteins in the mitochondrial fraction (Fig. 1B), the disappearance of these proteins treatment with chloramphenicol (CAP) (Fig. 1B), an inhibitor of prokaryotic/mitochondrial ribosomes (Grivell et al. 1971; Dunkle et al. 2010), and the assignment of 13 mtDNA-encoded proteins by their molecular weight (Supplemental Fig. S1D). Although AHA also enabled the detection of ANS-resistant signals, this analog provided a higher background signal (Supplemental Fig. S1E). The higher background signals may be due to the nonspecific labeling induced by terminal alkynes, as suggested in an earlier study (Speers and Cravatt 2004; Ali et al. 2019). Thus, in this study, we used HPG instead of AHA for the downstream assays.

**Figure 1.**
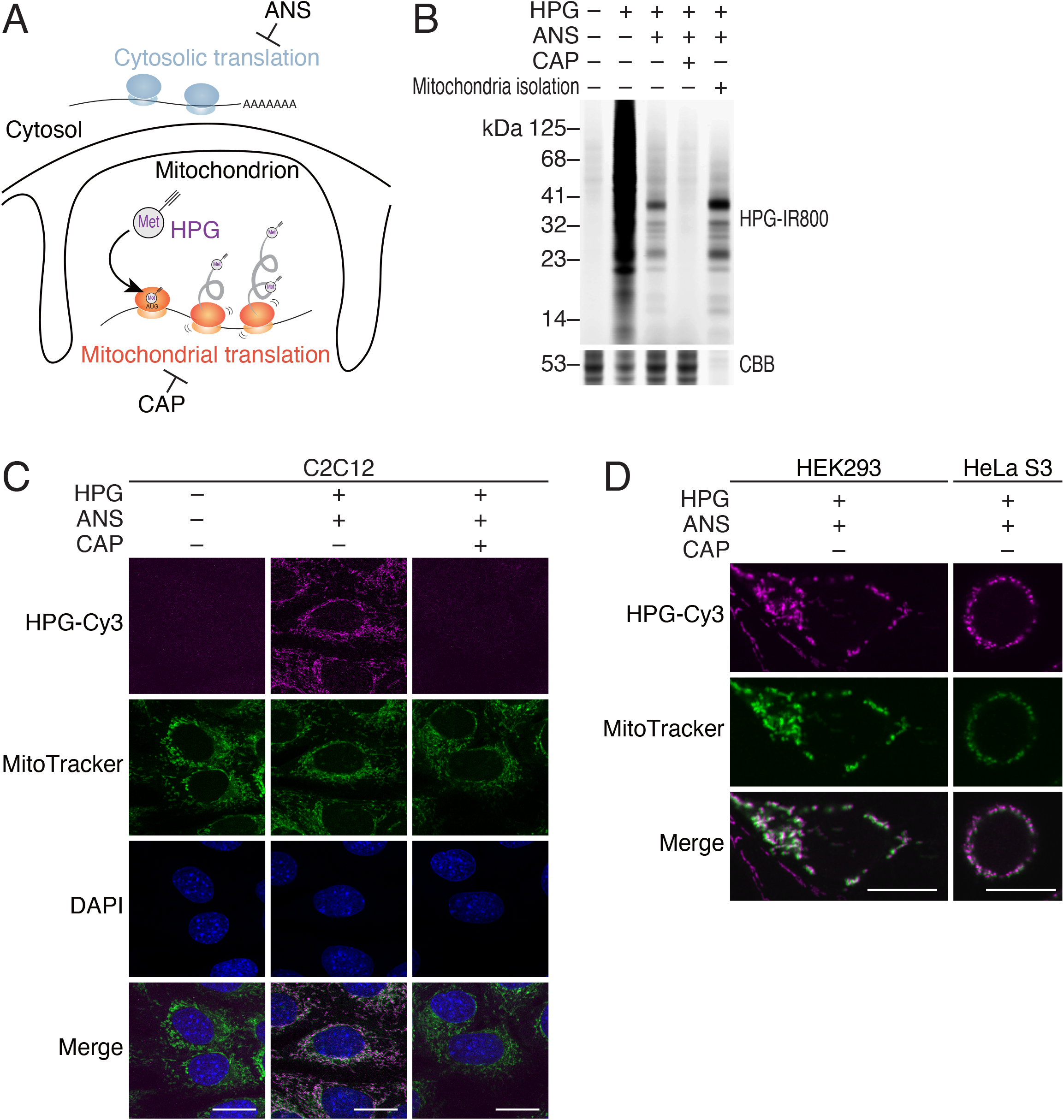
On-gel and *in situ* mito-FUNCAT in mammalian cell lines. (A) Schematic presentation of the experimental design. Mitochondrial nascent proteins were specifically labeled with a methionine analog, L-homopropargylglycine (HPG), while cytosolic translation was halted with anisomycin (ANS). As a control, mitochondrial translation was also blocked by chloramphenicol (CAP). (B) Representative gel images (n = 5) of total and mitochondrial nascent proteins. HEK293 cells were incubated with HPG in the presence or absence of inhibitors (ANS and CAP). Infrared (IR)-800 dye was conjugated to HPG-containing nascent proteins via a click reaction. Total protein was stained with CBB. (C) Representative microscopy images (n = 10) of mitochondrial nascent proteins *in situ*. Mouse C2C12 cells were incubated with HPG in the presence or absence of inhibitors (ANS and CAP). HPG-labeled proteins were visualized with Cy3. Mitochondria and nuclei were stained with MitoTracker Deep Red and DAPI, respectively. (D) Representative microscopy images of mitochondrial nascent proteins in the indicated human cell lines (HEK293, n = 9; HeLa S3, n = 4). HPG-labeled proteins were visualized with Cy3. Mitochondria were stained with MitoTracker. In C and D, the scale bar represents 15 μm.

Taking advantage of the lower background of HPG in the click reaction, we applied this technique to an in-cell reaction (*in situ* mito-FUNCAT) (Supplemental Fig. S1B). As expected, we observed that the FUNCAT signal was overlaid with a mitochondrial marker (MitoTracker) (Fig. 1C). This approach was not only restricted to mouse C2C12 cells (Fig. 1C), as reported in earlier work (Estell et al. 2017), but also succeeded in human HEK293 and HeLa S3 cells (Fig. 1D), suggesting the versatility of *in situ* mito-FUNCAT in a wide array of cells (Yousefi et al. 2021; Zorkau et al. 2021). Note that we used a lower concentration of HPG (50 μM) than earlier reports (Estell et al. 2017; Yousefi et al. 2021; Zorkau et al. 2021) (0.5-1 mM) to prevent the introduction of biases and/or artifacts with high doses of this compound. Under these conditions, we observed incremental accumulation of newly synthesized proteins in mitochondria over time (Supplemental Fig. S1F). Given the sufficiently high signal, we used a 3-h incubation condition throughout the experiments in this study.

### Implementation of mito-FUNCAT for FACS analysis

Next, we adapted the *in situ* mito-FUNCAT approach for use with flow cytometry (mito-FUNCAT-FACS) (Fig. 2A). Indeed, mito-FUNCAT-FACS on C2C12 cells (Supplemental Fig. 4A) successfully distinguished the signal of HPGylated mitochondrial nascent proteins (Fig. 2B +HPG +ANS) from the background (Fig. 2B – HPG). Reduction of the signal with CAP (Fig. 2B +HPG +ANS +CAP) further ensured the detection of mitochondrial translation products.

**Figure 2.**
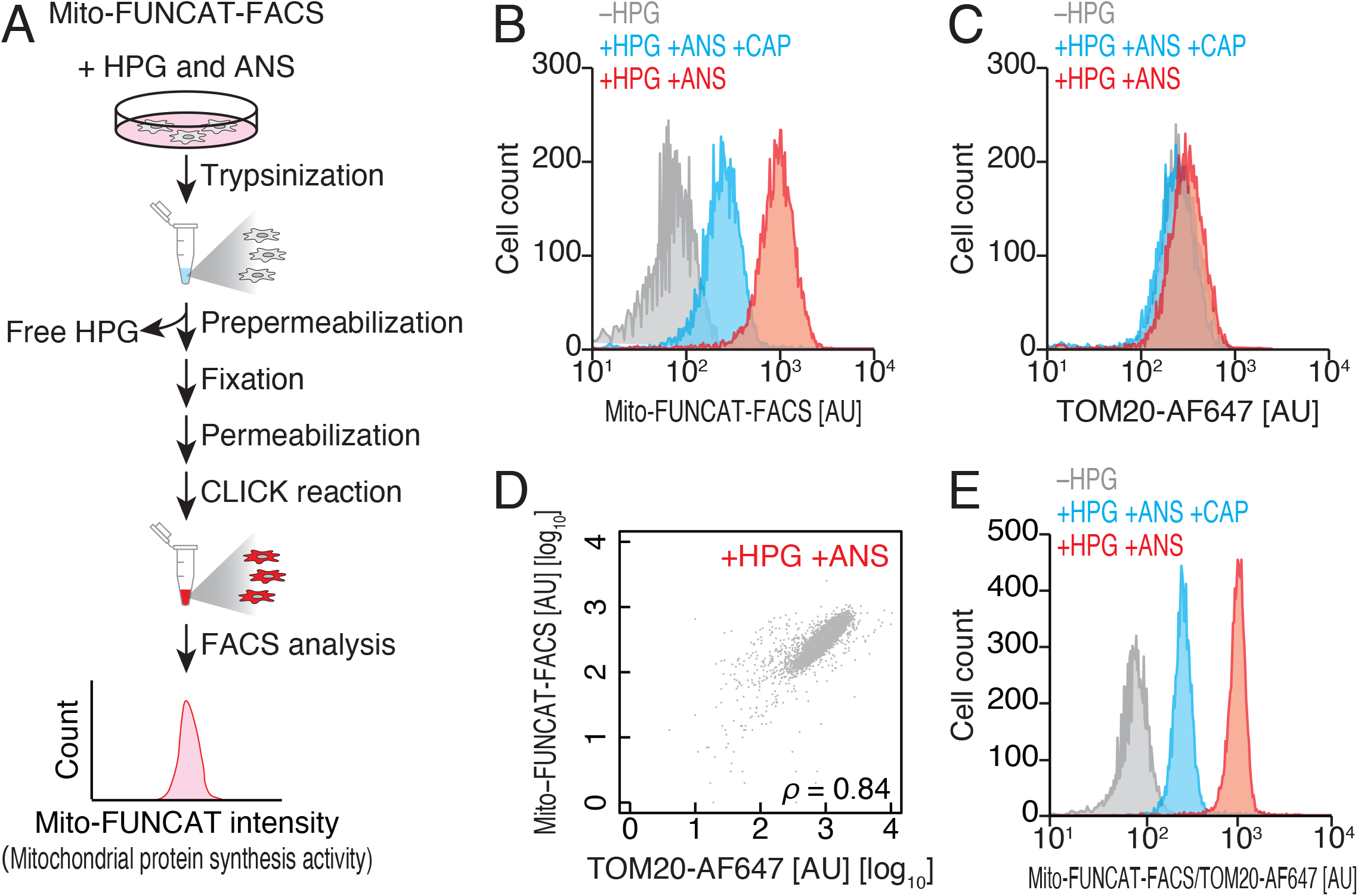
FACS-based quantification of mito-FUNCAT (mito-FUNCAT-FACS) (A) Schematic representation of the mito-FUNCAT-FACS procedure. (B and C) Representative distribution (n = 4) of Cy3-conjugated HPG signals (B, mito-FUNCAT-FACS) and Alexa Fluor (AF) 647-labeled TOM20 signals (C) across C2C12 cells in FACS. Cells were analyzed in the presence or absence of translation inhibitors (ANS and CAP) and HPG. Data from 1 × 10^4^ cells are shown. (D) Scatter plot for Cy3-conjugated HPG signals and AF647-labeled TOM20 signals across cells analyzed in B and C. ρ, Spearman’s rank correlation coefficient. (E) The distribution of mito-FUNCAT-FACS signals (in B) normalized to the AF647-labeled TOM20 intensity (in C). AU, arbitrary unit.

Since FACS allows us to track multiple fluorescent markers, we simultaneously measured mitochondrial abundance by translocase of outer mitochondrial membrane 20 (TOM20)—a receptor of presequence-carrying preproteins (Wiedemann and Pfanner 2017)—by immunostaining with a different fluorescent profile than the one used for the mitochondrial nascent peptides. This enabled the normalization of the mito-FUNCAT signal variation caused by differential mitochondrial masses in cells (Fig. 2C and 2D) and refined the resolution of mitochondrial translation measurements (Fig. 2E). Similar normalization could be conducted using MitoTracker Deep Red (Agnello et al. 2008; Cottet-Rousselle et al. 2011; Yeung et al. 2015) (Supplemental Fig. S2 and Supplemental Fig. 4B), although staining with dye depends on membrane potential.

### Mitochondrial function-associated translation activation monitored by mito-FUNCAT-FACS

To confirm the performance of mito-FUNCAT-FACS, we observed alterations in mitochondrial translation during organelle biogenesis. For this purpose, HeLa S3 cells were treated with bezafibrate, a peroxisome proliferator-activated receptor (PPAR) panagonist, which causes increased mitochondrial biogenesis and, thus, an increased number of OXPHOS complexes (Bastin et al. 2008; Wang and Moraes 2011; Wenz et al. 2011). An earlier work using L-[^35^S]-methionine metabolic labeling (Wenz et al. 2011) showed that bezafibrate may increase mitochondrial translation. Similarly, our on-gel mito-FUNCAT recapitulated the elevated mitochondrial protein synthesis with bezafibrate treatment (Fig. 3A). However, irrespective of the methods (L-[^35^S]-methionine or HPG), these data could not distinguish whether the enhanced protein synthesis originated from an increased abundance of mitochondria or increased translation in the organelle.

**Figure 3.**
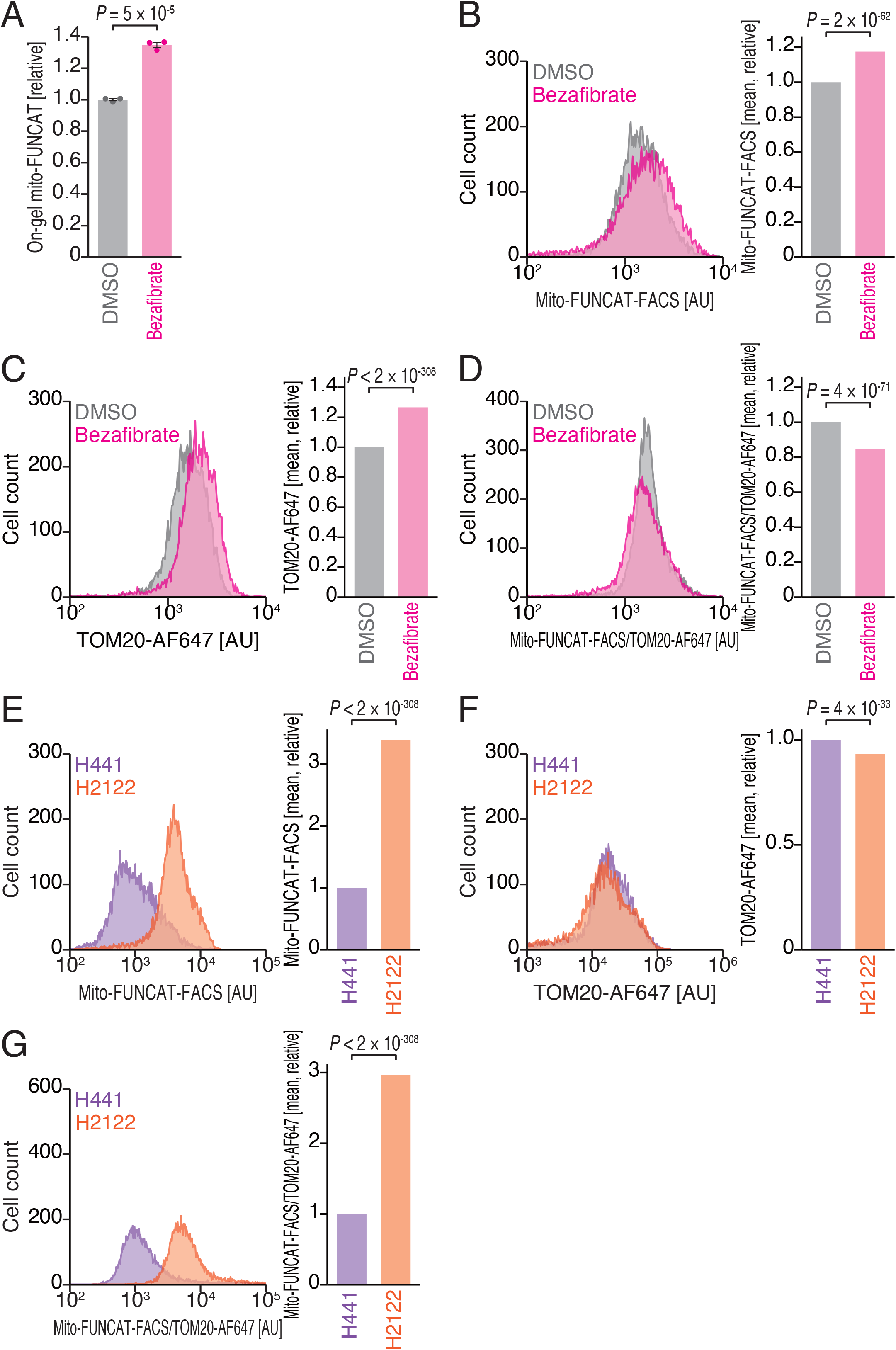
Mitochondrial protein synthesis under bezafibrate treatment assessed by mito-FUNCAT-FACS. (A) Bulk mitochondrial translation changes in HeLa S3 cells upon bezafibrate treatment, analyzed via on-gel mito-FUNCAT. Data from three replicates (points) and the mean (bar) with SD (error bar) are shown. The significance was calculated by Tukey’s test. (B and C) Representative distribution (n = 3) of Cy3-conjugated HPG signals (B, mito-FUNCAT-FACS) and AF647-labeled TOM20 signals (C) across HeLa S3 cells with or without bezafibrate treatment, analyzed by FACS. Data from 1 × 10^4^ cells are shown. The mean values (relative to the DMSO treatment) are shown in bar graphs. (D) The distribution of mito-FUNCAT-FACS signals (in B) normalized to the AF647-labeled TOM20 intensity (in C). The mean values (relative to DMSO treatment) are shown in bar graphs. (E-G) The same as B-D but comparing H441 and H2122 cells. Representative data (n = 4) are shown. In B-G, significance was calculated by the Mann–Whitney *U* test (two-tailed). AU, arbitrary unit.

Thus, we applied mito-FUNCAT-FACS and normalized the signals of HPGylated proteins according to the mitochondrial abundance measured via TOM20. The raw signals from HPGlyated nascent peptides showed an increase with bezafibrate treatment (Fig. 3B and Supplemental Fig. 4C), consistent with on-gel mito-FUNCAT (Fig. 3A). However, we observed a simultaneous increase in mitochondrial mass by TOM20 immunostaining (Fig. 3C). The normalization of mito-FUNCAT-FACS with the mitochondrial mass suppressed the scores (Fig. 3D), indicating that bezafibrate-mediated increase in HPGlyated proteins in mitochondria arose from the augmented organelle mass but not through a net increase in translation. Similar results were obtained from MitoTracker normalization (Supplemental Fig. S3 and Supplemental Fig. 4D). These data exemplified the ability of our FACS-based approach to disentangle the effects of mitochondrial translation and organelle biogenesis in individual cells.

To further evaluate the potential of mito-FUNCAT-FACS, we compared *in organello* translation in two different types of KRAS (Kirsten rat sarcoma viral oncogene homolog)-mutated lung cancer cell lines, H2122 and H441. H2122 cells possess mutations in the liver kinase B1 (LKB1) gene (also known as serine/threonine kinase 11 [STK11]) and Kelch-like ECH-associated protein 1 (KEAP1) gene, which are common mutations in lung adenocarcinoma, whereas H441 cells do not. It is commonly believed that the loss of LKB1 and KEAP1 leads to enhanced tricarboxylic acid (TCA) cycle flux (Faubert et al. 2014) and maintains high mitochondrial biosynthetic capacity (Kottakis et al. 2016). Consistent with the enhanced mitochondrial functions, the averaged signals of mito-FUNCAT-FACS in H2122 cells tended to be higher than those in H441 cells (Fig. 3E and Supplemental Fig. 4E). The increased mitochondrial translation occurred beyond the abundance of the organelle mass (Fig. 3F and 3G), suggesting that H2122 cells employed the increased mitochondrial translation to increase biosynthesis in the organelle.

### Mito-FUNCAT-FACS reveals heterogeneity of mitochondrial protein synthesis in cancer cells

In addition to utilizing averaged measurements, flow cytometry is useful for assessing cell subpopulations. We employed mito-FUNCAT-FACS to survey cellular heterogeneity in mitochondrial translation in various cell lines (C2C12, HeLa S3, HEK293T, A375, H441, H1944, H2009, and H2122) (Fig. 4A and Supplemental Fig. 4A-F). Indeed, we observed that H2009, H441, and H2122 cells showed a wide range of mitochondrial translation activity (Fig. 4A). The heterogeneity in H2122 cells was also evident from the *in situ* mito-FUNCAT, which highlighted a subset of cells exhibiting extremely high capacity of mitochondrial protein synthesis (Fig. 4B) (note that this *in situ* mito-FUNCAT was conducted with TOM20 immunostaining for the assessment of mitochondria, instead of MitoTracker staining, as shown in Fig. 1C and 1D). This phenotype was in contrast to C2C12 cells, which showed relatively uniform mitochondrial translation both in FACS and *in situ* microscopic options of mito-FUNCAT (Fig. 4A and 4B). The subpopulation of H2122 cells with high mitochondrial translation was associated with lower cell size and internal complexity, as indicated by forward and side scatter (FSC and SSC), suggesting unique characteristics of the cell population (Fig. 4C).

**Figure 4.**
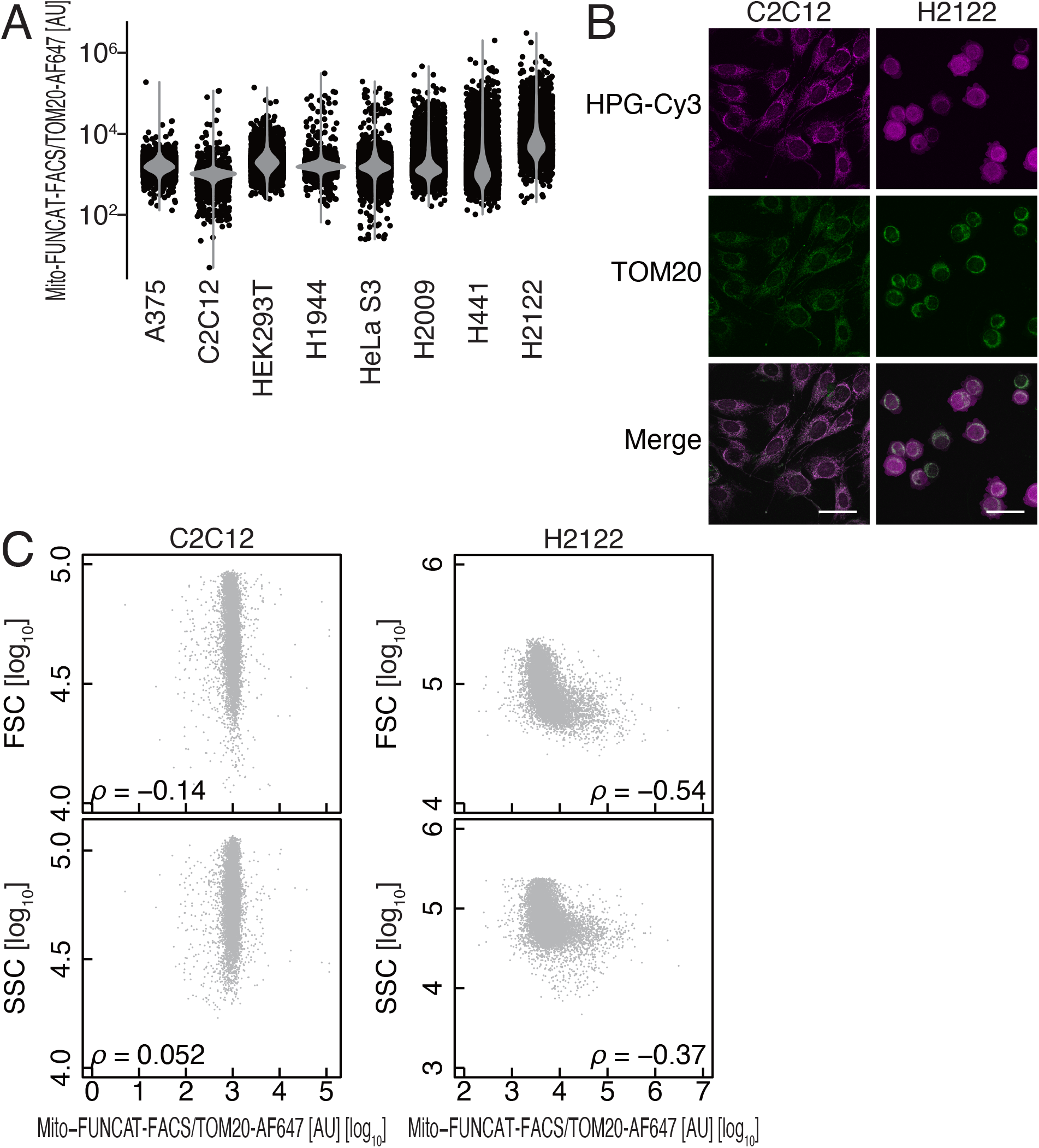
Heterogeneity of mitochondrial protein synthesis in H2122 cells. (A) Distributions of mito-FUNCAT-FACS scores (normalized to the AF647-labeled TOM20 intensity) in the indicated cell lines. Data from 1 × 10^4^ cells are shown. (B) Representative microscopy images (n = 2) of mitochondrial nascent proteins by by *in situ* mito-FUNCAT (Cy3-conjugated HPG. Mouse C2C12 and human H2122 cells were analyzed. Mitochondria were stained with anti-TOM20 antibodies. The scale bar represents 50 μm. (C) Scatter plots of mito-FUNCAT-FACS scores (TOM20-normalized) and forward scatter (FSC) or side scatter (SSC) in C2C12 and H2122 cells. ρ Spearman’s rank correlation coefficient. AU, arbitrary unit.

Thus, mito-FUNCAT-FACS developed in this work provides a useful framework for unveiling the previously-overlooked heterogeneity in mitochondrial protein synthesis among cell populations. These previously unreported differences in mitochondrial translation in H2122 cells (Fig. 4) needs further study to develop detailed characterizations. Given the ability of this technique to perform cell sorting and isolation, the assessment of gene expression profiles by bulk RNA-Seq and single-cell RNA-Seq can address the mechanism underlying variation in mitochondrial translation. Application of this experimental setup will pave the way for elucidating the *in organello* translation heterogeneity across a wide array of cell types, developmental stages, and responses to internal/external stimuli.

## Acknowledgments

We are grateful to Kosuke Dodo and all the members of the Iwasaki laboratory for constructive discussions, technical help, and critical reading of the manuscript. We also thank Kenji Ohtawa from the Support Unit for Bio-Material Analysis, RIKEN CBS Research Resources Division for FACS analysis and RIKEN CBS-Olympus Collaboration Center for performing the confocal microscopy. FACS analyses were also supported by Komaba Analysis Core, Institute of Industrial Science, The University of Tokyo. We also thank Shunsuke Kitajima for generous advice on mitochondrial functions in cancer cell lines. S.I. was supported by the Japan Society for the Promotion of Science (JSPS) (a Grant-in-Aid for Young Scientists [A], JP17H04998; a Challenging Research [Exploratory], JP19K22406), the Ministry of Education, Culture, Sports, Science and Technology (MEXT) (a Grant-in-Aid for Transformative Research Areas [B] “Parametric Translation”, JP20H05784), the Japan Agency for Medical Research and Development (AMED) (AMED-CREST, JP21gm1410001), and RIKEN (“Biology of Intracellular Environments” and “Integrated life science research to challenge super aging society”). Yoshiho I. was supported by JSPS (a Challenging Research [Pioneering], JP20K20643), MEXT (a Grant-in-Aid for Transformative Research Areas [B] “Parametric Translation”, JP20H05786), AMED (AMED-CREST, JP21gm1410001), and the Institute for AI and Beyond. T.O. was supported by JSPS (a Grant-in-Aid for Early-Career Scientists, JP20K20178), AMED (AMED-P-CREATE, G02-53), and the Takeda Science Foundation. Y.K. was supported by JSPS (a Grant-in-Aid for JSPS Fellows, JP20J10665). Y.K. was a RIKEN Junior Research Associate (JRA) Program recipient and a JSPS Research Fellow (DC2). H.S. was a RIKEN JRA Program recipient. T.W. was a recipient of the SPRING GX program from the University of Tokyo.

## Author contributions

Conceptualization, Y.K., H.S., T.O., and S.I.;

Methodology, Y.K., H.S., T.W., and S.I.;

Formal analysis, Y.K., H.S., T.O., Yasuhiro I., and T.W.;

Investigation, Y.K., H.S., T.O., Yasuhiro I., and T.W.;

Writing – Original Draft, Y.K. and S.I.;

Writing – Review & Editing, Y.K., H.S., T.O., Yasuhiro I., T.W., Yoshiho I., and S.I.;

Visualization, Y.K., H.S., T.O., T.W., and S.I.;

Supervision, Yoshiho I. and S.I.;

Project Administration, S.I.;

Funding Acquisition, Y.K., T.O., Yoshiho I., and S.I.

## Experimental procedures

### Cell culture

Cells were maintained in the following culture medium:

C2C12 (mouse myoblast, American Type Culture Collection [ATCC], CRL-1772), HeLa S3 (RIKEN BioResource Research Center, RCB1525), HEK293 (human embryonic kidney, ATCC, CRL-1573), HEK293T (ATCC, CRL-3216), A375 (human malignant melanoma, ATCC, CRL-1619), and H2009 (human lung adenocarcinoma, ATCC, CRL-5911) cells were maintained in DMEM, high glucose, GlutaMAX supplement (Thermo Fisher Scientific, 10566016) with 10% fetal bovine serum (FBS); H2122 (human lung adenocarcinoma, ATCC, CRL-5985), H1944 (human lung adenocarcinoma, ATCC, CRL-5907), and H441 (human lung adenocarcinoma, ATCC, HTB-174) cells were maintained in RPMI 1640 medium (Thermo Fisher Scientific, A1049101) with 10% FBS and 1% penicillin-streptomycin (Thermo Fisher Scientific, 15140148). All cells were cultured in a humidified incubator under 5% CO_2_ at 37°C.

The following compounds were used in this study: ANS (Chem-Impex International), CAP (FUJIFILM Wako Pure Chemical Corporation), CHX (Sigma-Aldrich), EME (Cayman Chemical), and bezafibrate (FUJIFILM Wako Pure Chemical Corporation).

### On-gel mito-FUNCAT

Cells were washed with PBS and incubated in methionine-free DMEM with 50 μM HPG (Jena Bioscience) and 100 μg/mL ANS, CHX, or EME for 3 h. For mitochondrial translation inhibition, 100 μg/mL CAP was added to the medium. Then, the cells were washed with ice-cold PBS and lysed with lysis buffer (20 mM Tris-HCl pH 7.5, 150 mM NaCl, 5 mM MgCl_2_, and 1% Triton X-100). The lysate was cleared by centrifugation at 20,000 *g* at 4 °C for 10 min. The supernatant was used for the click reaction with 50 μM azide-IRDye800CW (LI-COR Biosciences) by a Click-iT Cell Reaction Buffer Kit (Thermo Fisher Scientific) according to the manufacturer’s instructions, followed by SDS-PAGE. Images of labeled nascent peptides were acquired with an Odyssey CLx Infrared Imaging System (LI-COR Biosciences) in the 800 nm channel. Subsequently, the gel was stained with Coomassie brilliant blue (CBB) using EzStain AQua (ATTO) and imaged in the 700 nm channel to measure the protein input. The images were quantified with Image Studio version 5.2 (LI-COR Biosciences).

For mitochondrial isolation, EzSubcell Fraction (ATTO) was used according to the manufacturer’s instructions. The purified mitochondria were lysed with lysis buffer.

For AHA-mediated mito-FUNCAT, AHA (Thermo Fisher Scientific) and alkyne-IRDye800CW (LI-COR Biosciences) were used instead of HPG and azide-IRDye800CW, respectively.

### *In situ* mito-FUNCAT

Typically, 2 × 10^4^ cells were cultured on a laminin-coated Lab-Tek II Chamber Slide (Thermo Fisher Scientific) in 200 μL of methionine-free DMEM with HPG and ANS as described in the *On-gel mito-FUNCAT* section. For mitochondrial translation inhibition, 100 μg/mL CAP was added to the medium. Cells were washed with 200 μL of pre-warmed PBS, pre-permeabilized with 0.0005% digitonin in 200 μL of mitochondrial protective buffer (10 mM HEPES-KOH pH 7.5, 300 mM sucrose, 10 mM NaCl, and 5 mM MgCl_2_) for 5 min, and fixed in 200 μL of 4% paraformaldehyde (PFA) for 15 min. Then, the cells were fully permeabilized with 0.1% (v/v) Triton X-100 in 200 μL of PBS for 5 min and subsequently incubated in 100 μL of click reaction buffer (1× Click-iT cell reaction buffer [Thermo Fisher Scientific], 2 mM CuSO_4_, 1× Click-iT cell buffer additive [Thermo Fisher Scientific], and 1 μM azide-conjugated Cy3 [(Jena Bioscience]) for 30 min according to the Click-iT Cell Reaction Buffer Kit (Thermo Fisher Scientific) to label nascent proteins via the click reaction. After washing with 200 μL of Intercept Blocking Buffer (LI-COR Biosciences), mitochondria were immunostained in 100 μL of Intercept Blocking Buffer containing 1 μL of Alexa Fluor 647 Anti-TOM20 antibody (Abcam, ab209606) for 1 h at 4°C. The cells were washed with 200 μL of PBS twice, and then the nuclei were stained with 4′,6-diamidino-2-phenylindole (DAPI) (Thermo Fisher Scientific) for 5 min. Images were obtained using an FV3000 confocal microscope (Olympus) with a 60× objective lens (Olympus Japan, UPLSAPO60XS2).

For C2C12 and HeLa S3 cells, mitochondria were stained with 100 nM MitoTracker Deep Red FM (Thermo Fisher Scientific) for 15 min during HPG labeling, omitting TOM20 immunostaining.

Images were colored by standard look-up tables (LUTs) in FV3000 software and exported in TIF format (24 bit). ImageJ2 (https://github.com/imagej/imagej2, version 2.3.0) was used to overlay images.

### Mito-FUNCAT-FACS

Cells were handled as described in the *In situ mito-FUNCAT* section with some modifications. Approximately 1 × 10^6^ cells were cultured in a 10-cm dish with 10 mL of methionine-free DMEM with HPG and ANS as described in the *On-gel mito-FUNCAT* section. Then, the cells were washed with 5 mL of PBS, trypsinized with 1.5 mL of 0.05% trypsin-EDTA (Thermo Fisher Scientific), collected into 2.0-mL tubes, and washed with 500 μL of PBS. Pre-permeabilization, fixation, permeabilization, click reaction, and immunostaining were conducted in a 2.0-mL tube as described in *In situ mito-FUNCAT*, using 500 μL of mitochondrial protective buffer with 0.0005% digitonin, 500 μL of 4% PFA, 500 μL of 0.1% (v/v) Triton X-100 in PBS, 250 μL of click reaction buffer, and 100 μL of Intercept Blocking Buffer containing 1 μL of Alexa Fluor 647 Anti-TOM20 antibody, respectively. Cells were collected by centrifugation at 300 *g* for 3 min before fixation and by centrifugation at 1,000 *g* for 3 min after fixation. Cells were analyzed with a FACSAria II cell sorter or FACSMelody (BD Biosciences). Data from 1 × 10^4^ cells are depicted in the graphs.

---

**Supplemental Figure 1.**
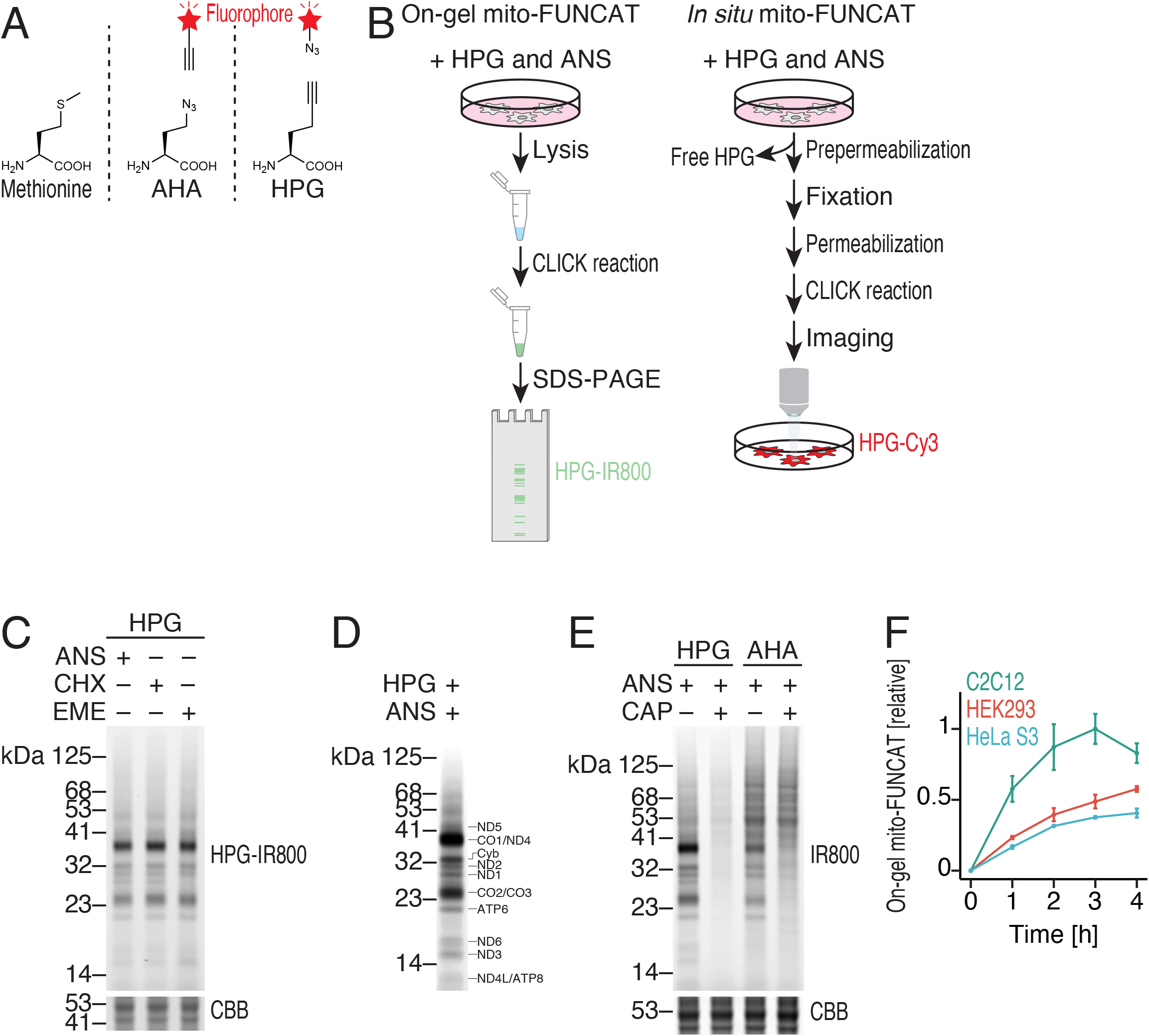
Characterization of newly synthesized proteins in mitochondria. (A) Methionine analogs and click chemistry-compatible ligands used for metabolic labeling of mitochondrial nascent proteins. (B) Schematic representations of on-gel mito-FUNCAT and *in situ* mito-FUNCAT procedures. (C) Representative gel images (n = 3) of total and mitochondrial nascent proteins. HEK293 cells were incubated with HPG in the presence of cytosolic translation inhibitors, ANS, CHX, or EME. Infrared (IR)-800 dye was conjugated to HPG-containing nascent proteins via a click reaction. Total protein was stained with CBB. (D) The assignment of 13 mitochondrial genome-encoded proteins to proteins detected with on-gel mito-FUNCAT. We note that because of its hydrophobic nature, CO1, which was predicted to be 57 kDa, aberrantly migrated around the 37-kDa region. (E) Mitochondrial nascent proteins (n = 2) were labeled with AHA or HPG and with azide-IR800 or alkyne-IR800, respectively. CAP was used to block mitochondrial translation and to monitor the background signal. (F) Time course experiments of on-gel mito-FUNCAT for the indicated cell lines. The mean value after 3-h incubation in C2C12 cells was set to 1. The mean (point) with SD (error bar) from three replicates are shown.

**Supplemental Figure 2.**
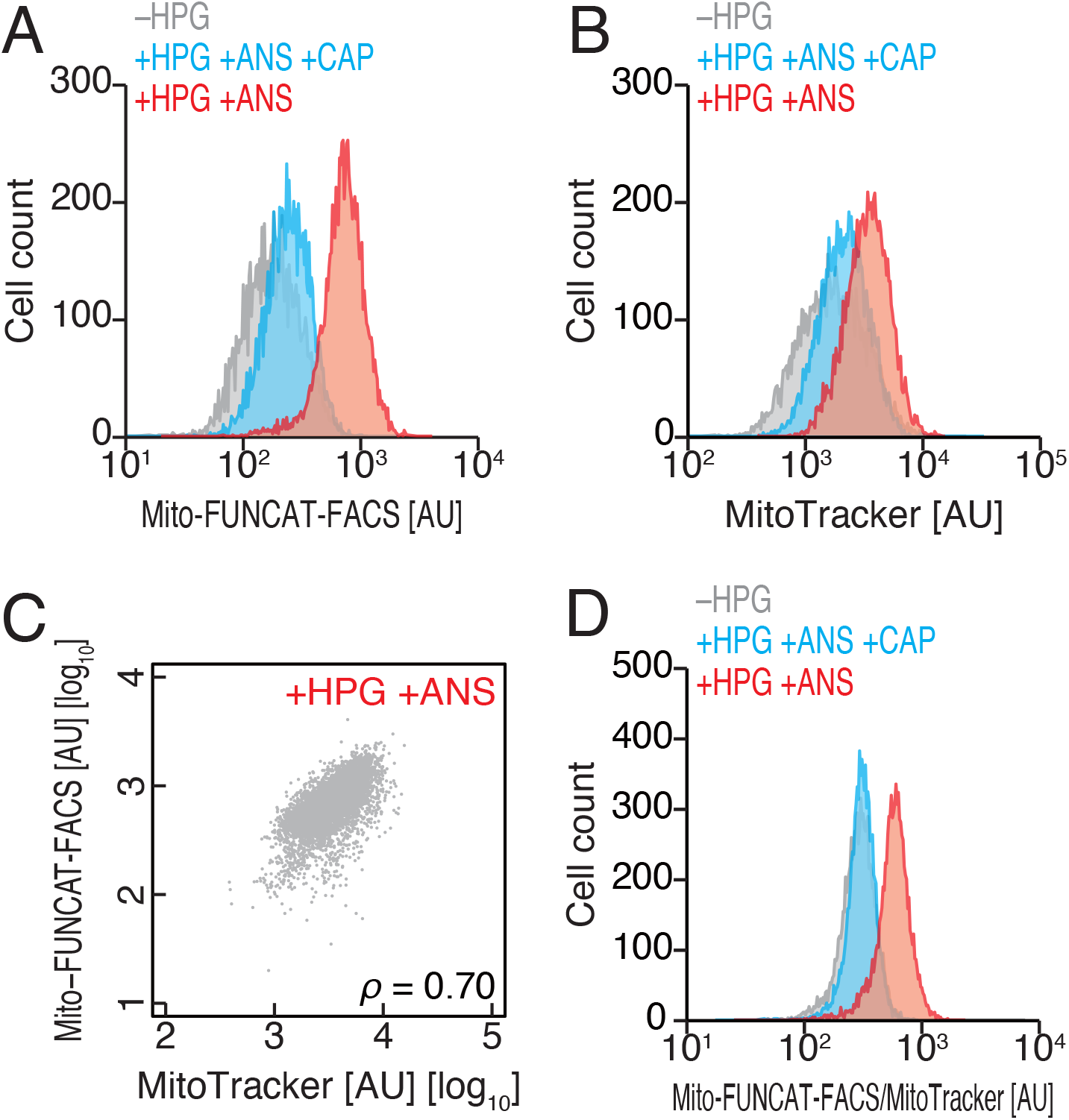
Mito-FUNCAT-FACS signals normalized by MitoTracker. (A-D) The same experiments as in Fig. 2B-E but the mitochondria mass was measured and normalized by MitoTracker Deep Red. Data from 1 × 10^4^ cells are shown. Representative data (n = 5) are shown.

**Supplemental Figure 3.**
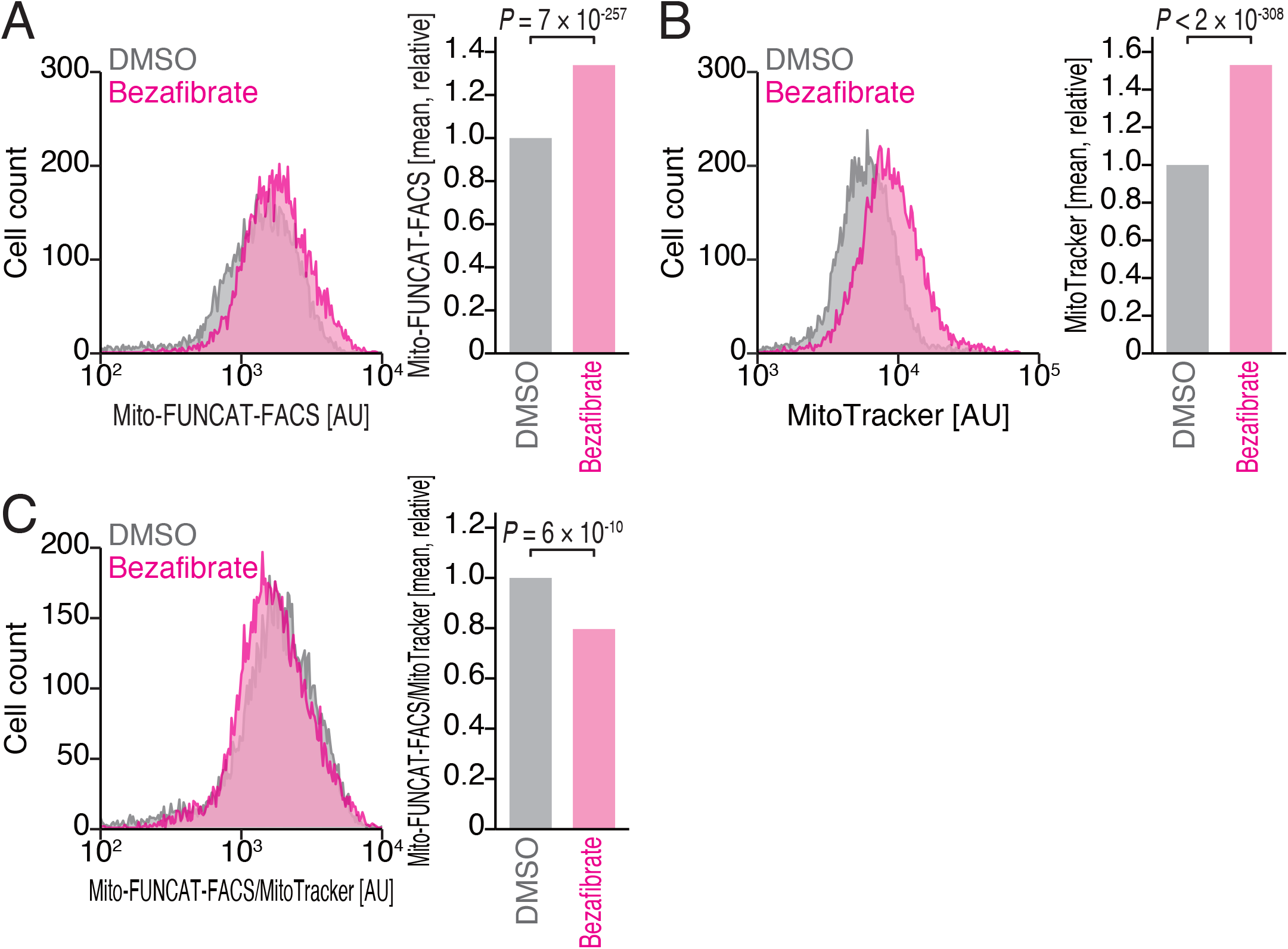
Normalization of mito-FUNCAT-FACS signals in bezafibrate treatment by MitoTracker. (A-C) The same experiments as in Fig. 3B-D but the mitochondria mass was measured and normalized by MitoTracker Deep Red. Data from 1 × 10^4^ cells are shown. Representative data (n = 2) are shown.

**Supplemental Figure 4.**
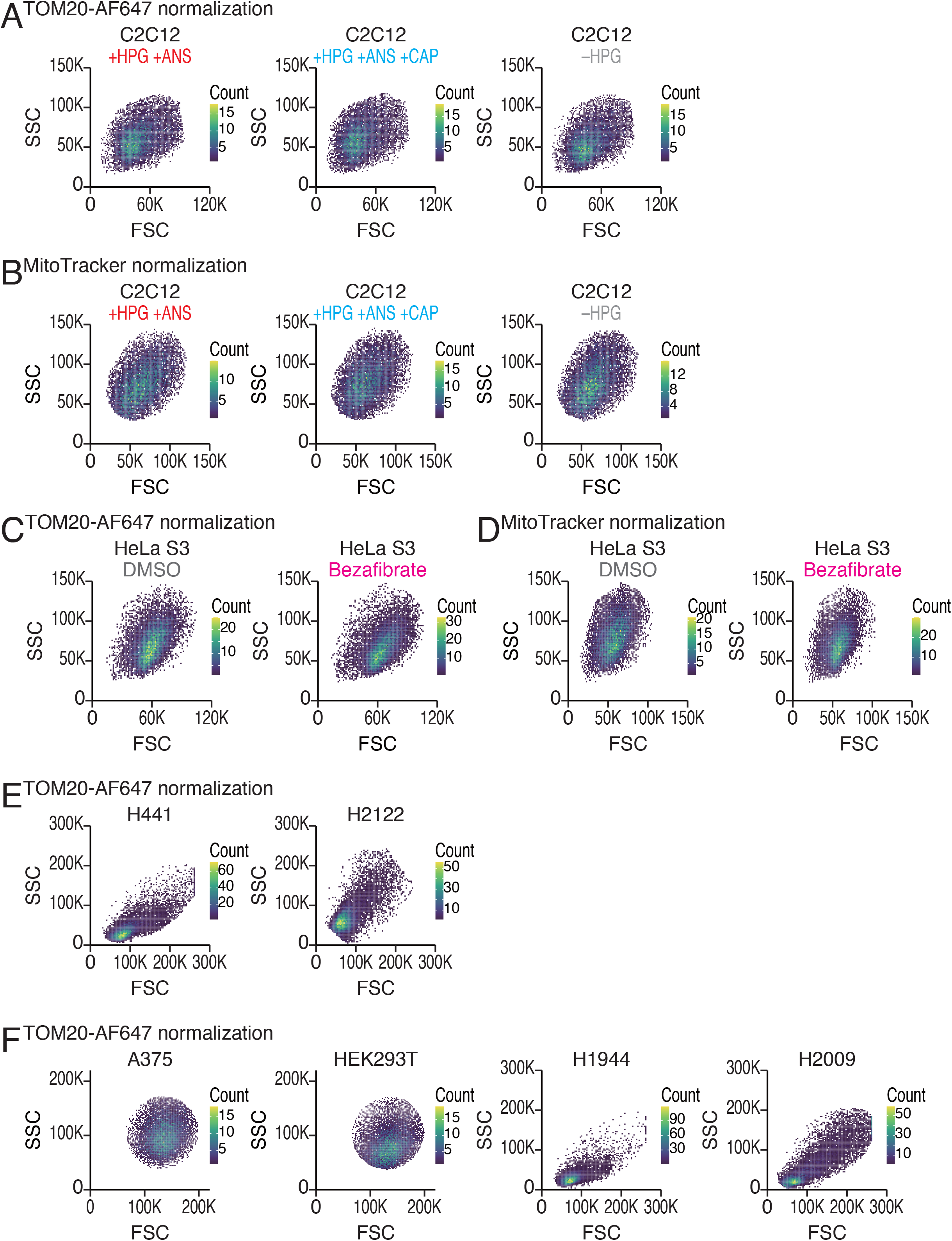
Characterization of gated cells in mito-FUNCAT-FACS. (A-F) Distributions of forward scatter (FSC) and side scatter (SSC) of cells analyzed in mito-FUNCAT-FACS (in Figs. 2-4 and Supplemental Figs. S2-3).

## Notes

### Competing Interest Statement

The authors have declared no competing interest.

## References

1. Agnello M, Morici G, Rinaldi AM. 2008. A method for measuring mitochondrial mass and activity. Cytotechnology 56: 145–149.

2. Ali TH, Heidelberg T, Hussen RSD, Tajuddin HA. 2019. Unexpected reactions of terminal alkynes in targeted “Click chemistry” coppercatalyzed azide-alkyne cycloadditions. Curr Org Synth 16: 1143–1148.

3. Anderson S, Bankier AT, Barrell BG, de Bruijn MH, Coulson AR, Drouin J, Eperon IC, Nierlich DP, Roe BA, Sanger F, et al. 1981. Sequence and organization of the human mitochondrial genome. Nature 290: 457–465.

4. Antonicka H, Sasarman F, Nishimura T, Paupe V, Shoubridge EA. 2013. The mitochondrial RNA-binding protein GRSF1 localizes to RNA granules and is required for posttranscriptional mitochondrial gene expression. Cell Metab 17: 386–398.

5. Asano K, Suzuki T, Saito A, Wei F-Y, Ikeuchi Y, Numata T, Tanaka R, Yamane Y, Yamamoto T, Goto T, et al. 2018. Metabolic and chemical regulation of tRNA modification associated with taurine deficiency and human disease. Nucleic Acids Res 46: 1565–1583.

6. Bastin J, Aubey F, Rötig A, Munnich A, Djouadi F. 2008. Activation of peroxisome proliferator-activated receptor pathway stimulates the mitochondrial respiratory chain and can correct deficiencies in patients’ cells lacking its components. J Clin Endocrinol Metab 93: 1433–1441.

7. Beatty KE, Liu JC, Xie F, Dieterich DC, Schuman EM, Wang Q, Tirrell DA. 2006. Fluorescence visualization of newly synthesized proteins in mammalian cells. Angew Chem Int Ed Engl 45: 7364–7367.

8. Beatty KE, Tirrell DA. 2008. Two-color labeling of temporally defined protein populations in mammalian cells. Bioorg Med Chem Lett 18: 5995–5999.

9. Cahoon AB, Qureshi AA. 2018. Leaderless mRNAs are circularized in *Chlamydomonas reinhardtii* mitochondria. Curr Genet 64: 1321–1333.

10. Cottet-Rousselle C, Ronot X, Leverve X, Mayol JF. 2011. Cytometric assessment of mitochondria using fluorescent probes. Cytometry A 79: 405–425.

11. Couvillion MT, Churchman LS. 2017. Mitochondrial ribosome (mitoribosome) profiling for monitoring mitochondrial translation in vivo. Curr Protoc Mol Biol 119: 4.28.1–4.28.25.

12. Couvillion MT, Soto IC, Shipkovenska G, Churchman LS. 2016. Synchronized mitochondrial and cytosolic translation programs. Nature 533: 499–503.

13. Cruz-Zaragoza LD, Dennerlein S, Linden A, Yousefi R, Lavdovskaia E, Aich A, Falk RR, Gomkale R, Schöndorf T, Bohnsack MT, et al. 2021. An *in vitro* system to silence mitochondrial gene expression. Cell 184: 5824–5837.e15.

14. De Silva D, Tu Y-T, Amunts A, Fontanesi F, Barrientos A. 2015. Mitochondrial ribosome assembly in health and disease. Cell Cycle 14: 2226–2250.

15. Dieterich DC, Hodas JJ, Gouzer G, Shadrin IY, Ngo JT, Triller A, Tirrell DA, Schuman EM. 2010. *In situ* visualization and dynamics of newly synthesized proteins in rat hippocampal neurons. Nat Neurosci 13: 897–905.

16. Dunkle JA, Xiong L, Mankin AS, Cate JH. 2010. Structures of the *Escherichia coli* ribosome with antibiotics bound near the peptidyl transferase center explain spectra of drug action. Proc Natl Acad Sci U S A 107: 17152–17157.

17. Englmeier R, Pfeffer S, Förster F. 2017. Structure of the human mitochondrial ribosome studied *in situ* by cryoelectron tomography. Structure 25: 1574–1581.e2.

18. Estell C, Stamatidou E, El-Messeiry S, Hamilton A. 2017. *In situ* imaging of mitochondrial translation shows weak correlation with nucleoid DNA intensity and no suppression during mitosis. J Cell Sci 130: 4193–4199.

19. Faubert B, Vincent EE, Griss T, Samborska B, Izreig S, Svensson RU, Mamer OA, Avizonis D, Shackelford DB, Shaw RJ, et al. 2014. Loss of the tumor suppressor LKB1 promotes metabolic reprogramming of cancer cells via HIF-1α. Proc Natl Acad Sci U S A 111: 2554–2559.

20. Fung S, Nishimura T, Sasarman F, Shoubridge EA. 2013. The conserved interaction of C7orf30 with MRPL14 promotes biogenesis of the mitochondrial large ribosomal subunit and mitochondrial translation. Mol Biol Cell 24: 184–193.

21. Gao F, Wesolowska M, Agami R, Rooijers K, Loayza-Puch F, Lawless C, Lightowlers RN, Chrzanowska-Lightowlers ZMA. 2017. Using mitoribosomal profiling to investigate human mitochondrial translation. Wellcome Open Res 2: 116.

22. Garreau de Loubresse N, Prokhorova I, Holtkamp W, Rodnina MV, Yusupova G, Yusupov M. 2014. Structural basis for the inhibition of the eukaryotic ribosome. Nature 513: 517–522.

23. Grimes BT, Sisay AK, Carroll HD, Cahoon AB. 2014. Deep sequencing of the tobacco mitochondrial transcriptome reveals expressed ORFs and numerous editing sites outside coding regions. BMC Genomics 15: 31.

24. Grivell LA, Reijnders L, de Vries H. 1971. Altered mitochondrial ribosomes in a cytoplasmic mutant of yeast. FEBS Lett 16: 159–163.

25. Hinz FI, Dieterich DC, Tirrell DA, Schuman EM. 2012. Non-canonical amino acid labeling in vivo to visualize and affinity purify newly synthesized proteins in larval zebrafish. ACS Chem Neurosci 3: 40–49.

26. Imami K, Selbach M, Ishihama Y. 2021. Monitoring mitochondrial translation by pulse SILAC. bioRxiv. https://www.biorxiv.org/content/10.1101/2021.01.31.428997v1.abstract.

27. Ingolia NT, Ghaemmaghami S, Newman JR, Weissman JS. 2009. Genome-wide analysis *in vivo* of translation with nucleotide resolution using ribosome profiling. Science 324: 218–223.

28. Iwasaki S, Floor SN, Ingolia NT. 2016. Rocaglates convert DEAD-box protein eIF4A into a sequence-selective translational repressor. Nature 534: 558–561.

29. Iwasaki S, Ingolia NT. 2017. The growing toolbox for protein synthesis studies. Trends Biochem Sci 42: 612–624.

30. Jeffreys AJ, Craig IW. 1976. Analysis of proteins synthesized in mitochondria of cultured mammalian cells. An assessment of current approaches and problems in interpretation. Eur J Biochem 68: 301–311.

31. Kashiwagi K, Shichino Y, Osaki T, Sakamoto A, Nishimoto M, Takahashi M, Mito M, Weber F, Ikeuchi Y, Iwasaki S, et al. 2021. eIF2B-capturing viral protein NSs suppresses the integrated stress response. Nat Commun 12: 1–12.

32. Keilland E, Rupar CA, Prasad AN, Tay KY, Downie A, Prasad C. 2016. The expanding phenotype of MELAS caused by the m.3291T > C mutation in the *MT-TL1* gene. Mol Genet Metab Rep 6: 64–69.

33. Kirino Y, Goto Y-I, Campos Y, Arenas J, Suzuki T. 2005. Specific correlation between the wobble modification deficiency in mutant tRNAs and the clinical features of a human mitochondrial disease. Proc Natl Acad Sci U S A 102: 7127–7132.

34. Kottakis F, Nicolay BN, Roumane A, Karnik R, Gu H, Nagle JM, Boukhali M, Hayward MC, Li YY, Chen T, et al. 2016. LKB1 loss links serine metabolism to DNA methylation and tumorigenesis. Nature 539: 390–395.

35. Kummer E, Ban N. 2021. Mechanisms and regulation of protein synthesis in mitochondria. Nat Rev Mol Cell Biol 22: 307–325.

36. Lee M, Matsunaga N, Akabane S, Yasuda I, Ueda T, Takeuchi-Tomita N. 2021. Reconstitution of mammalian mitochondrial translation system capable of correct initiation and long polypeptide synthesis from leaderless mRNA. Nucleic Acids Res 49: 371–382.

37. Li SH-J, Nofal M, Parsons LR, Rabinowitz JD, Gitai Z. 2021. Monitoring mammalian mitochondrial translation with MitoRiboSeq. Nat Protoc 16: 2802–2825.

38. Morscher RJ, Ducker GS, Li SH-J, Mayer JA, Gitai Z, Sperl W, Rabinowitz JD. 2018. Mitochondrial translation requires folate-dependent tRNA methylation. Nature 554: 128–132.

39. Pearce SF, Rorbach J, Van Haute L, D’Souza AR, Rebelo-Guiomar P, Powell CA, Brierley I, Firth AE, Minczuk M. 2017. Maturation of selected human mitochondrial tRNAs requires deadenylation. Elife 6: e27596.

40. Pfeffer S, Woellhaf MW, Herrmann JM, Förster F. 2015. Organization of the mitochondrial translation machinery studied *in situ* by cryoelectron tomography. Nat Commun 6: 6019.

41. Roche FK, Marsick BM, Letourneau PC. 2009. Protein synthesis in distal axons is not required for growth cone responses to guidance cues. J Neurosci 29: 638–652.

42. Rooijers K, Loayza-Puch F, Nijtmans LG, Agami R. 2013. Ribosome profiling reveals features of normal and disease-associated mitochondrial translation. Nat Commun 4: 2886.

43. Rötig A. 2011. Human diseases with impaired mitochondrial protein synthesis. Biochim Biophys Acta 1807: 1198–1205.

44. Scharfe C, Lu HH-S, Neuenburg JK, Allen EA, Li G-C, Klopstock T, Cowan TM, Enns GM, Davis RW. 2009. Mapping gene associations in human mitochondria using clinical disease phenotypes. PLoS Comput Biol 5: e1000374.

45. Schöller E, Marks J, Marchand V, Bruckmann A, Powell CA, Reichold M, Mutti CD, Dettmer K, Feederle R, Hüttelmaier S, et al. 2021. Balancing of mitochondrial translation through METTL8-mediated m^3^C modification of mitochondrial tRNAs. Mol Cell 81: 4810–4825.e12.

46. Signer RA, Magee JA, Salic A, Morrison SJ. 2014. Haematopoietic stem cells require a highly regulated protein synthesis rate. Nature 509: 49–54.

47. Speers AE, Cravatt BF. 2004. Profiling enzyme activities in vivo using click chemistry methods. Chem Biol 11: 535–546.

48. Suomalainen A, Battersby BJ. 2018. Mitochondrial diseases: the contribution of organelle stress responses to pathology. Nat Rev Mol Cell Biol 19: 77–92.

49. Suzuki T, Yashiro Y, Kikuchi I, Ishigami Y, Saito H, Matsuzawa I, Okada S, Mito M, Iwasaki S, Ma D, et al. 2020. Complete chemical structures of human mitochondrial tRNAs. Nat Commun 11: 4269.

50. Tcherkezian J, Brittis PA, Thomas F, Roux PP, Flanagan JG. 2010. Transmembrane receptor DCC associates with protein synthesis machinery and regulates translation. Cell 141: 632–644.

51. Wang X, Moraes CT. 2011. Increases in mitochondrial biogenesis impair carcinogenesis at multiple levels. Mol Oncol 5: 399–409.

52. Webb BD, Diaz GA, Prasun P. 2020. Mitochondrial translation defects and human disease. J Transl Genet Genom 4: 71–80.

53. Wenz T, Wang X, Marini M, Moraes CT. 2011. A metabolic shift induced by a PPAR panagonist markedly reduces the effects of pathogenic mitochondrial tRNA mutations. J Cell Mol Med 15: 2317–2325.

54. Weraarpachai W, Antonicka H, Sasarman F, Seeger J, Schrank B, Kolesar JE, Lochmüller H, Chevrette M, Kaufman BA, Horvath R, et al. 2009. Mutation in *TACO1*, encoding a translational activator of COX I, results in cytochrome c oxidase deficiency and late-onset Leigh syndrome. Nat Genet 41: 833–837.

55. Wiedemann N, Pfanner N. 2017. Mitochondrial machineries for protein import and assembly. Annu Rev Biochem 86: 685–714.

56. Wong W, Bai X-C, Brown A, Fernandez IS, Hanssen E, Condron M, Tan YH, Baum J, Scheres SHW. 2014. Cryo-EM structure of the *Plasmodium falciparum* 80S ribosome bound to the anti-protozoan drug emetine. Elife 3: e03080.

57. Yasukawa T, Suzuki T, Ishii N, Ueda T, Ohta S, Watanabe K. 2000a. Defect in modification at the anticodon wobble nucleotide of mitochondrial tRNA^Lys^ with the MERRF encephalomyopathy pathogenic mutation. FEBS Lett 467: 175–178.

58. Yasukawa T, Suzuki T, Suzuki T, Ueda T, Ohta S, Watanabe K. 2000b. Modification defect at anticodon wobble nucleotide of mitochondrial tRNAs^Leu^(UUR) with pathogenic mutations of mitochondrial myopathy, encephalopathy, lactic acidosis, and stroke-like episodes. J Biol Chem 275: 4251–4257.

59. Yeung M, Hurren R, Nemr C, Wang X, Hershenfeld S, Gronda M, Liyanage S, Wu Y, Augustine J, Lee EA, et al. 2015. Mitochondrial DNA damage by bleomycin induces AML cell death. Apoptosis 20: 811–820.

60. Yoon BC, Jung H, Dwivedy A, O’Hare CM, Zivraj KH, Holt CE. 2012. Local translation of extranuclear lamin B promotes axon maintenance. Cell 148: 752– 764.

61. Yousefi R, Fornasiero EF, Cyganek L, Montoya J, Jakobs S, Rizzoli SO, Rehling P, Pacheu-Grau D. 2021. Monitoring mitochondrial translation in living cells. EMBO Rep 22: e51635.

62. Zhang X, Zuo X, Yang B, Li Z, Xue Y, Zhou Y, Huang J, Zhao X, Zhou J, Yan Y, et al. 2014. MicroRNA directly enhances mitochondrial translation during muscle differentiation. Cell 158: 607–619.

63. Zorkau M, Albus CA, Berlinguer-Palmini R, Chrzanowska-Lightowlers ZMA, Lightowlers RN. 2021. High-resolution imaging reveals compartmentalization of mitochondrial protein synthesis in cultured human cells. Proc Natl Acad Sci U S A 118: e2008778118.

